# An intuitive method to visualize Boolean network attractors landscape based on probability distribution from trajectories

**DOI:** 10.1101/2024.10.31.621222

**Authors:** Ket Hing Chong

**Affiliations:** School of Computer Science and Engineering, Nanyang Technological University, Singapore 639798

**Keywords:** Boolean network attractors landscape, directed graph, probability distribution, Waddington’s landscape, data science application

## Abstract

**Motivation:** Boolean network modeling is a coarse-grained modeling approach that has been highly valued for its simple abstract representation of biological regulation in molecular network. The simplified ‘On’ and ‘Off’ expression of gene or protein product is a qualitative representation of experimental findings for gene gets turned on or turned off. Boolean network constructed can be analyzed as having point attractor or cyclic attractor; however, current softwares can visualize the Boolean network attractors on a plane and not visualizing Boolean attractors’ landscape in a 3 dimensional view.

**Results:** Here, we propose an intuitive method to visualize Boolean network attractors landscape based on probability distribution *P* of all possible trajectories in the state transition graph. We formulate a quasi-potential value *U* = − *ln P* that enables us to plot the Boolean network attractors landscape in 3D. There are two types of attractors in Boolean network: cyclic attractor and point attractor. Our intuitive method can visualize Boolean network in a landscape with point attractor (node labeled with yellow color) and cyclic attractor (node labeled with pink color). In the 3D landscape the attractors are located at the bottom of the directed graphs. The propose method can be applied in the study of networks in general, for example applications in directed graph, social network for popular users or e-commerce store to find favorite products as identified by attractors.

**Availability and implementation:** The source code of Python implementation of the propose method can be downloaded from Boolean-net-attractor-Land URL link.

## 1. Introduction

Boolean network modeling is an attractive coarse-grain method to model gene regulatory network using discrete logical description^1,2,3^. The logical Boolean network modeling framework is assumed to be an approximation of certain genes get turned ‘On’ or turned ‘Off’ by the qualitative representation of 1 or 0^4,5,6^. It has been widely used in modeling gene regulatory network to gain insights on the dynamical behavior of biological system^7,8^, and became a powerful tool for analyzing large gene regulatory network^9,10^. Recently, Boolean network attractor landscape analysis has been used to study DNA damage response in p53 network feedback loops^8^ and colorectal tumorigenesis by modeling of human signaling network^11^. Many softwares have been developed to facilitate modeling and analysis of Boolean gene regulatory networks, such as GINsim^12,13^, BoolNet^14,15^ and Attractor Landscape Analysis Toolbox (ATLANTIS)^16^. GINsim and ATLANTIS use graphical user interface design. BoolNet is a command line software package in R. However, GINsim and BoolNet can plot Boolean network in a plane to visualize the state transition graphs and attractors (point or cyclic attractor). Cyclic attractor is also known as cycle in a directed graph. For instance cyclic attractor represents the p53 oscillations in a cell state for cell cycle arrest^8^. Whereas point attractor is used to represents a stable cell state for cell senescence or apoptosis^16^. One of the main difficulties (in the case of GINsim and BoolNet) is the 2D visualization of the Boolean attractors landscape. Whereas ATLANTIS requires the use of MATLAB software (R2016a) to run the toolbox for plotting the 3D landscape for Boolean Network. There are a few works published in the literature on visualizing Boolean network landscape^17,8^, however, these methods are for probabilistic Boolean network.

Here, we propose an intuitive method for visualizing deterministic Boolean network attractors landscape in 3D view. The method is adapted from our previous Monte Carlo method for quantifying Waddington’s epigenetic landscape for continuous variables in ordinary differential equations (ODEs)^18,19^.

## 2. Material and methods

In order to plot Waddington’s epigenetic landscape we require to find a quasi-potential *U* by estimating the steady state probability distribution *P*. The quasi-potential value is given by *U* = − *lnP* ^20,21,22,18^. We adopted the idea of Monte Carlo framework from our previous work on quantifying Waddington’s epigenetic landscape^18^ for continuous variables to estimate the probability distribution (*P*) from large number of trajectories simulated from random initial conditions. In a similar idea, we estimate probability distribution from all trajectories in the state transition graph in Boolean network. Based on all the trajectories from state transition graph we can estimate a probability distribution. For example, in a trajectory [2, 1, 0] where it starts from node 2, then goes to node 1 and ends at node 0. Because this trajectory passes node 1 and node 0, we can add 1 to the counting variable named **countn** to node 1 and node 0. From the counting **countn** calculate the total as the sum of all the nodes. Then, we can estimate a probability distribution by dividing each element in **countn** with the sum.

To demonstrate the intuitive method we created a toy model of Boolean network shown in Figure 1. The toy model consists of 15 nodes. There are two point attractors (node 5 and node 11). One cyclic attractor forms by node 0 and node 1. In our intuitive method for plotting the landscape for Boolean network we need to list all the possible trajectories for each node. Then based on all the possible trajectories for each node we can calculate the probability distribution *P*. We formulate a quasi-potential value *U* = − *ln P* that enables us to plot the Boolean network attractors landscape in 3D. We use Python to implement the calculation of probability distribution and drawing of the 3D landscape for Boolean network.

**Figure 1:**
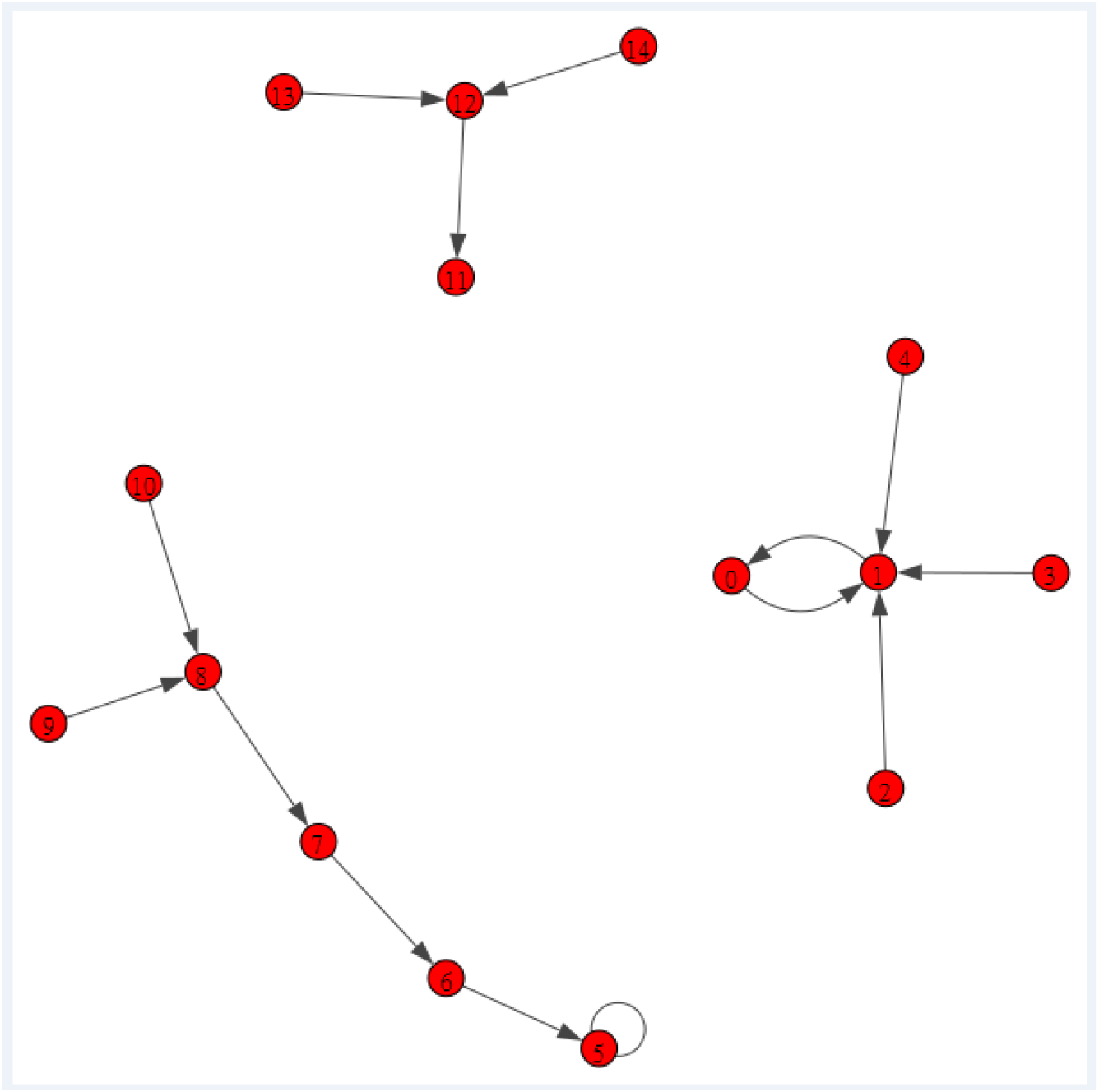
Figure above shows a toy model of Boolean gene regulatory network. We use this toy model to demonstrate the intuitive method for drawing the landscape of Boolean network. There are 15 nodes in the Boolean network. Node 5 and node 11 are point attractors. Node 0 and node 1 forms a cyclic attractor.

## 3. Results

To demonstrate our intuitive method to plot Boolean network attractors landscape we selected four models as case studies. The first model is a toy model to demonstrate the intuitive method. The second model is a simple model from GINsim. The third model is a model of cell cycle by Fauré *et al*. (2006)^23^. The fourth model is a social network model from Facebook NIPS 2012^24^.

### 3.1 Case Study 1: Toy model

For our toy model, we can list all the possible trajectories with no repeated node in the Boolean network or directed graph. The possible trajectories to each node are given in Table 1. From all the possible trajectories to each node we can count the number of trajectory to each node. To avoid zero probability we added 1 to each count for the number of trajectory to each node. Then, we can obtain the probability distribution *P* as shown in Table 2.

**Table 1.** The possible trajectories to each node where trajectory cannot have repeated node except self loop (or self directed node).

| node | no. of trajectory | Trajectory |
| --- | --- | --- |
| 0 | 4 | [1, 0] ; [2, 1, 0] ; [3, 1, 0] ; [4, 1, 0] |
| 1 | 4 | [0, 1] ; [2, 1] ; [3, 1] ; [4, 1] |
| 2 | 0 | nil |
| 3 | 0 | nil |
| 4 | 0 | nil |
| 5 | 6 | [5, 5] ; [6, 5] ; [7, 6, 5] ; [8, 7, 6, 5] ; [9, 8, 7, 6, 5] ; [10, 8, 7, 6, 5] |
| 6 | 4 | [7, 6] ; [8, 7, 6] ; [9, 8, 7, 6] ; [10, 8, 7, 6] |
| 7 | 3 | [8, 7] ; [9, 8, 7] ; [10, 8, 7] |
| 8 | 2 | [9, 8] ; [10, 8] |
| 9 | 0 | nil |
| 10 | 0 | nil |
| 11 | 3 | [12, 11] ; [13, 12, 11] ; [14, 12, 11] |
| 12 | 2 | [13, 12] ; [14, 12] |
| 13 | 0 | nil |
| 14 | 0 | nil |
| total | 28 |  |

**Table 2.** The probability distribution in the toy model of Boolean network. It is calculated based on the number of trajectory to each node. To avoid zero probability value, we added 1 to each node.

| node | no. of trajectory | no. of trajectory plus 1 | probability P |
| --- | --- | --- | --- |
| 0 | 4 | 5 | 5/43 |
| 1 | 4 | 5 | 5/43 |
| 2 | 0 | 1 | 1/43 |
| 3 | 0 | 1 | 1/43 |
| 4 | 0 | 1 | 1/43 |
| 5 | 6 | 7 | 7/43 |
| 6 | 4 | 5 | 5/43 |
| 7 | 3 | 4 | 4/43 |
| 8 | 2 | 3 | 3/43 |
| 9 | 0 | 1 | 1/43 |
| 10 | 0 | 1 | 1/43 |
| 11 | 3 | 4 | 4/43 |
| 12 | 2 | 3 | 3/43 |
| 13 | 0 | 1 | 1/43 |
| 14 | 0 | 1 | 1/43 |
| total | 28 | 43 | 1 |

However, in our implementation we replace the add one to each node by adding its own node. Since, the self-loop just contributed one trajectory and we considered it is not important for probability distribution so we ignore the self-loop. Another reason is we consider trajectories with no repeated node. The results from our implementation are given below.

The trajectories to each node are:

node [0]= [[0], [1, 0], [2, 1, 0], [3, 1, 0], [4, 1, 0]]

node [1]= [[0, 1], [1], [2, 1], [3, 1], [4, 1]]

node [2]= [[2]]

node [3]= [[3]]

node [4]= [[4]]

node [5]= [[5], [6, 5], [7, 6, 5], [8, 7, 6, 5], [9, 8, 7, 6, 5], [10, 8, 7, 6, 5]]

node [6]= [[6], [7, 6], [8, 7, 6], [9, 8, 7, 6], [10, 8, 7, 6]]

node [7]= [[7], [8, 7], [9, 8, 7], [10, 8, 7]]

node [8]= [[8], [9, 8], [10, 8]]

node [9]= [[9]]

node [10]= [[10]]

node [11]= [[11], [12, 11], [13, 12, 11], [14, 12, 11]]

node [12]= [[12], [13, 12], [14, 12]]

node [13]= [[13]]

node [14]= [[14]].

The total number of trajectories is 42. Therefore, the probability distribution is almost the same as calculated manually 43 as in Table 2.

Based on this intuitive probability distribution *P* we can calculate the potential *U* for each node by using the formula U=-ln P ^20,21,22,18^. The 3D landscape for the toy model is shown in Figure 2. The 3D landscape for Boolean network shows the attractors (point attractor and cyclic attractor) located at the bottom of the landscape.

**Figure 2:**
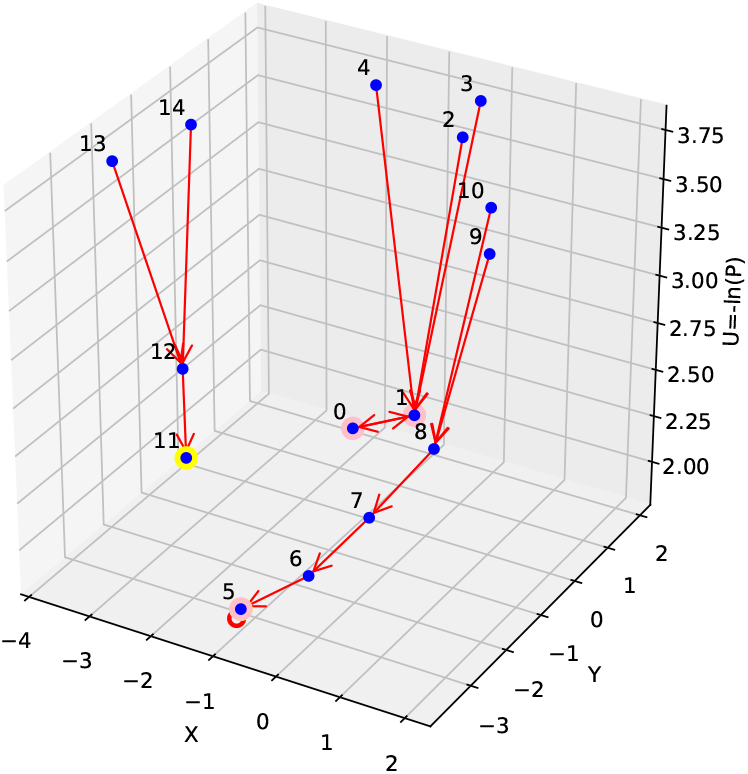
The demonstration of the intuitive method using a toy model of Boolean gene regulatory network. The 3D landscape of the toy model of Boolean network plotted with our method. Based on Figure 1 the directed graph we know node 11 is point attractor. Whereas node 0 and node 1 forms a cyclic attractor. Node 5 has a self loop and can be classified as cyclic attractor as well.

After testing on the Toy model, we check the accuracy of the implementation with a slightly more connected network (Example 1 in Tutorial Igraph with Python) proposed by Chire ^25^. We validated the accuracy of the method with this model and the results are provided in the Supplementary Information.

### 3.2 Case Study 2: Example model from GINsim

For the second case study, we selected an example of Boolean network from Gene Interaction Network simulation (GINsim) software^12^. GINsim is a software tool for creating and simulation of Boolean Network model. In this model there are three genes: G1, G2 and G3. The state transition graph of this Boolean network model is given in Figure 3. The state transition graph is drawn in 2 dimension plane. In GINsim it allows user to select the different layout views such as Level, Ring, and 3D. The state transition graph generated by GINsim with the 3D layout is given in Figure 3. In this model there is one point attractor (node labeled with 200) and the rest of the nodes are cyclic attractors. The 3D landscape plotted with our intuitive method is shown in Figure 4. The 3D landscape of Boolean network indicates the point attractor node 200 labeled with yellow color is located at the bottom and the rest of the cyclic attractors labeled with pink color. The difference between 3D view layout in GINsim and 3D landscape from our method is the node plotted from our method indicates all stable attractors such as point attractor or cyclic attractor are located at the bottom. It is more intuitive and follows the concept of Waddington’s landscape to illustrates stable states or attractors.

**Figure 3:**
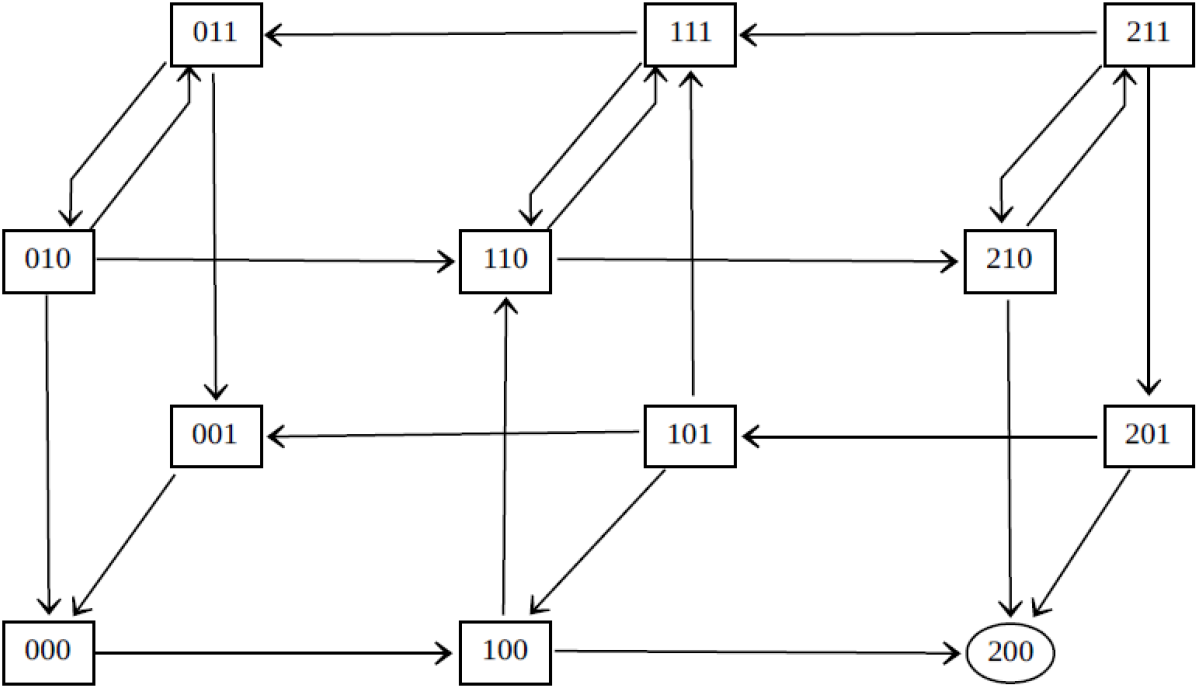
The state transition graph of a Boolean network model example from GINsim is given above. This state transition graph is generated by using GINsim. There is one point attractor: 200. The rest of the nodes are cyclic attractors.

**Figure 4:**
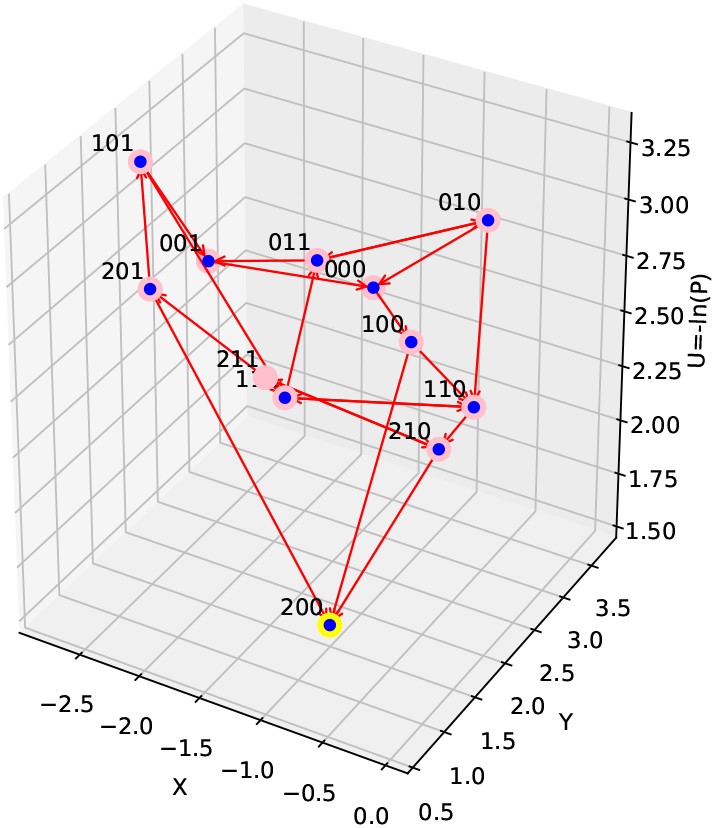
The 3D landscape of the model of Boolean network example from GINsim plotted with our method. Based on Figure 3 the state transition graph we know node 200 is a point attractor (node labeled yellow). Whereas the other nodes forms cyclic attractors (node labeled pink).

### 3.3 Case Study 3: cell cycle model by Fauré *et al*. (2006) from BoolNet

For the third case study, we selected a biological network in cell cycle ^23^. This cell cycle model is created by BoolNet. The cell cycle network contains 10 genes: CycD, Rb, E2F, CycE, CycA, p27, Cdc20, Cdh1, UbcH10 and CycB. To run BoolNet it requires the igraph package installed. BoolNet is a command line package in R with a documentation given in BoolNet Package Vignette^14^. From BoolNet we run the cell cycle model and we obtain the state transition graph as given in Figure 5. In the state transition graph we identified two basins of attractions. The attractor 1 labeled with blue color consists of a group of nodes converging to the node 162 for [0100010100]; 1 means the gene is active and 0 means the gene is not active. The attractor 2 labeled with green color consists of another group of nodes converging to the cyclic attractors formed by 7 nodes: [1001100000], [1000100011], [1000101011], [1000001110], [1010000110], [1011000100], [1011100100].

**Figure 5:**
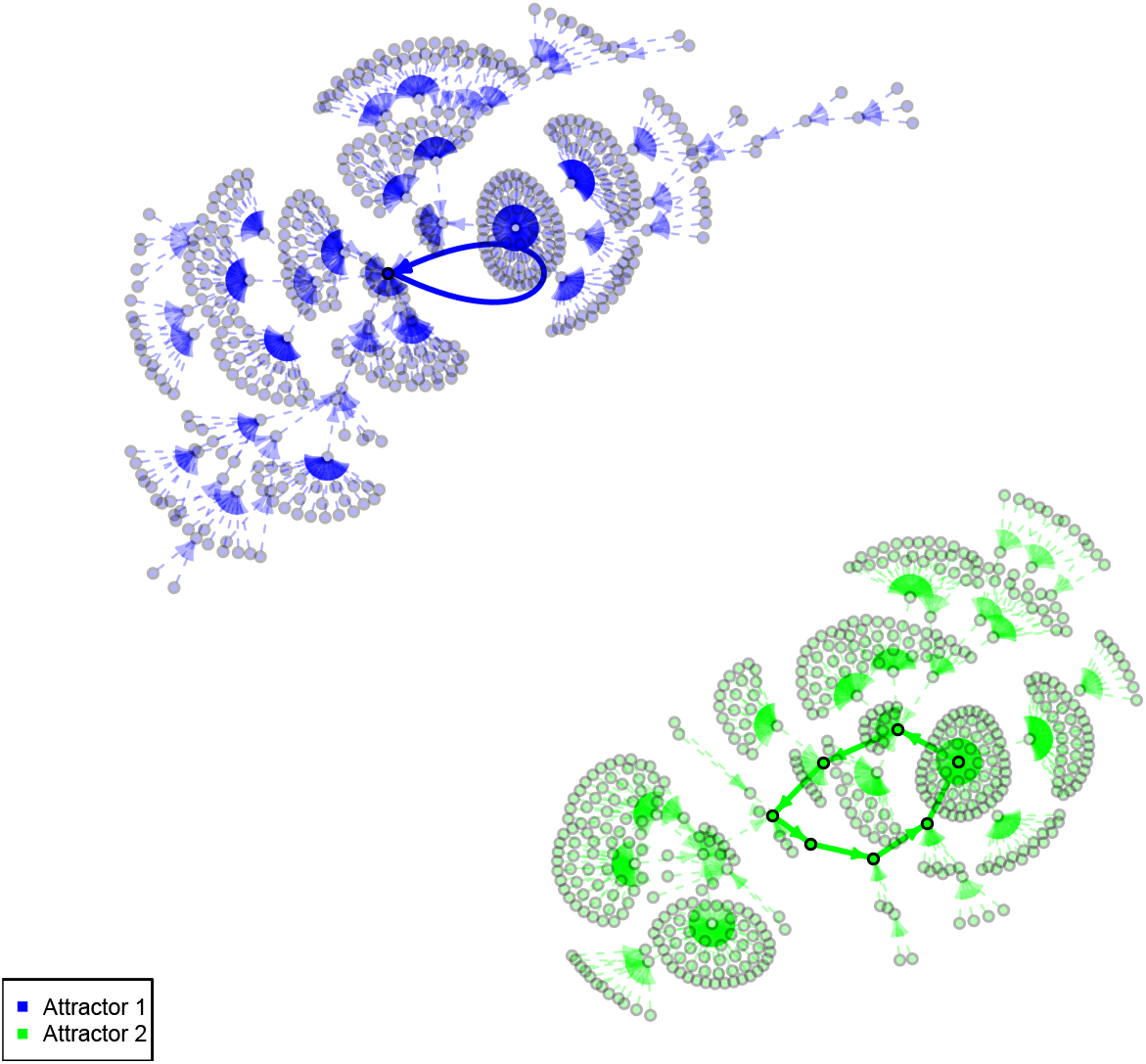
The state transition graph of Boolean gene regulatory network for cell cycle model^23^ drawn by using BoolNet. This state transition graph in BoolNet can visually display the two basins of attraction. Attractor 1 consists of the nodes labeled with blue color and attractor 2 represents the nodes labeled with green color. In attractor 1 there is a self-loop. Whereas in attractor 2 there is a cyclic attractor (7 nodes drawn with black circle).

From BoolNet we exported the model information to a .net file called cellcycle.net. With the model information we can apply our intuitive method for plotting the 3D landscape of this network attractors. The result is shown in Figure 6. Same as the result from BoolNet, there are two basins of attraction. Attractor 1 is formed by a group of nodes on the left of Figure 6. Attractor 2 is formed by another group of nodes on the right of Figure 6. The results are equivalent only with a more intuitive description of the landscape in 3D.

**Figure 6:**
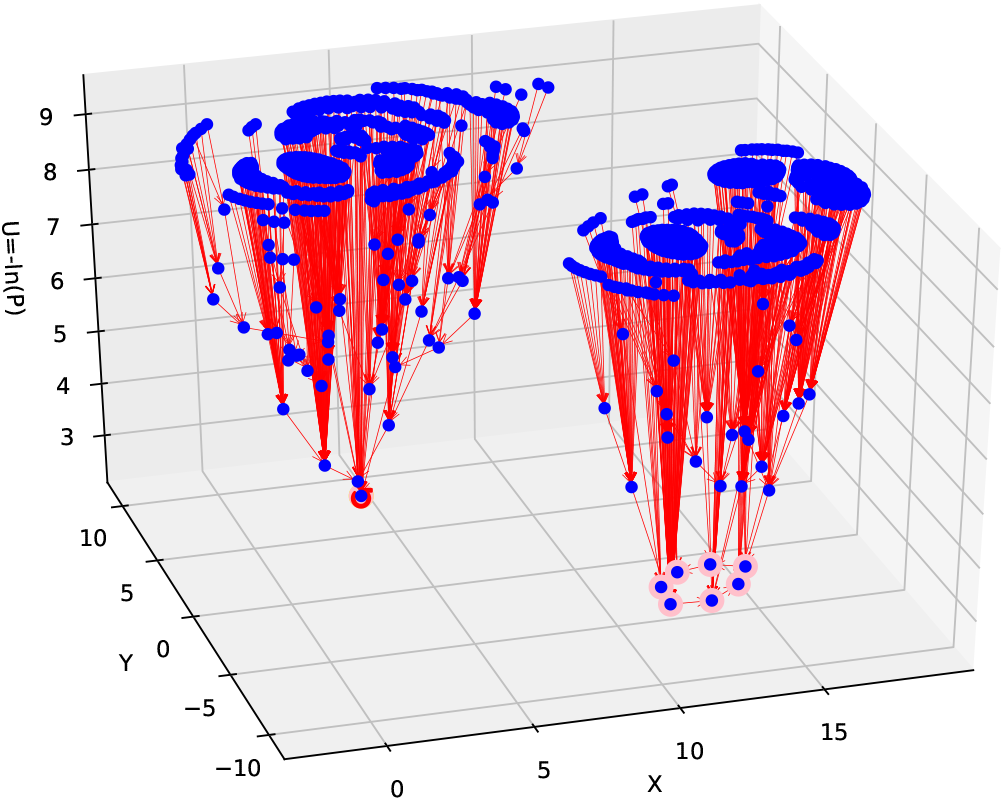
The demonstration of the intuitive method using a model of Boolean gene regulatory network for cell cycle^23^. The 3D landscape of the cell cycle model plotted with our method. There are two groups of attractors. The first group is attractor 1 on the left (same as the blue color nodes in Figure 5), there is a self-loop in node 162. The second group is on the right (same as the green color nodes in Figure 5). Attractor 2 at the bottom is cyclic attractors formed by the 7 nodes [25, 785, 849, 449, 389, 141, 157] (node labeled pink).

### 3.4 Case Study 4: social network model from Facebook NIPS 2012

For the fourth case study, we selected a social network from Facebook NIPS 2012. Facebook NIPS data were downloaded from the KONECT Project^24^. The nodes represent users. The edges represent friendships where node from the arrow is a friend to another user (node in which arrow point to). In this network there are 2,888 users and 2,981 friendships. The 3D landscape generated from our method is shown in Figure 7. From Figure 7 we can see there are 3 point attractors for user number: 1, 2536 and 2687. An attractor is a user or node as stable point attractor. Social network is known to be large and complex network. There is no good method to organize social network^26^. Here, we demonstrated the propose method for quantifying and plotting of Boolean network can also be applied to social network to visualize and identified the users as point attractors. Where point attractor can be interpreted as popular user who is having great influence in the social network from Facebook NIPS 2012 data.

**Figure 7:**
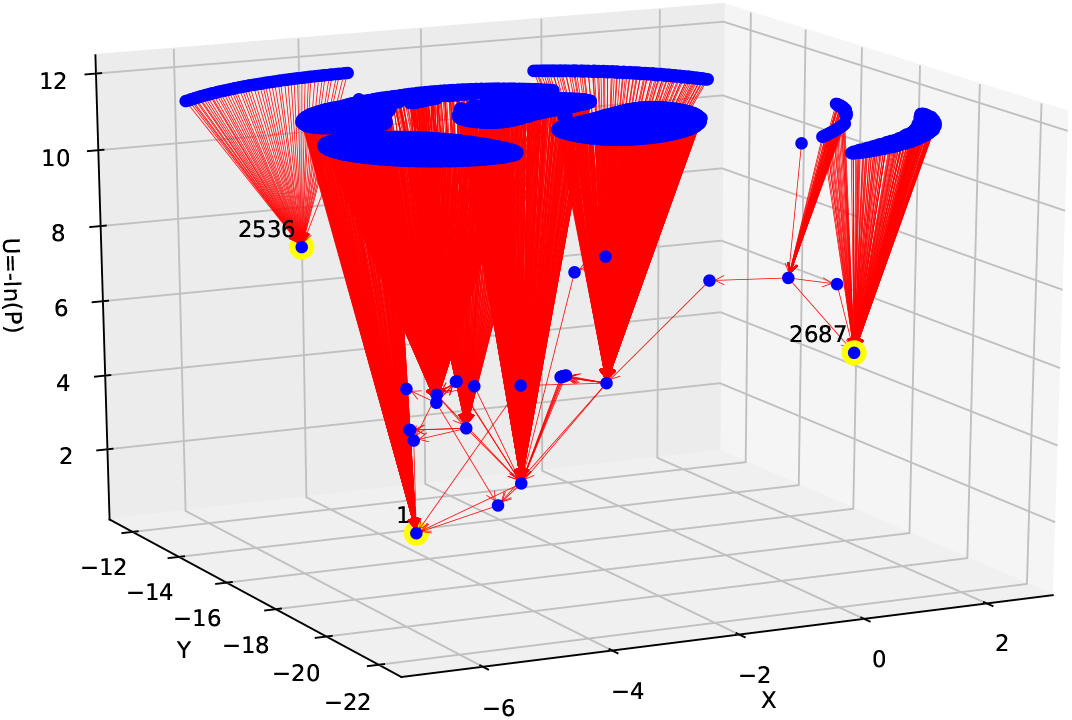
The 3D landscape of a social network model from Facebook NIPS 2012 is given above. There are 3 point attractors (node labeled yellow) for user number 1, 2536 and 2687. There are no cyclic attractors.

## 4. Conclusion

In this work we present a method to plot 3D landscape for Boolean network attractors. This intuitive method is adapted from our previous work of Monte Carlo method for ODEs^18^. From the four case studies above we have explained and demonstrated the usefulness of the method. The propose method in general can be used for application in data science and the study of networks such as directed graph, social network for popular users or e-commerce store to identified favorite products as characterized by attractors.

## Credit authorship contribution statement

### Ket Hing Chong

Designed the method, Implemented the algorithm, Case studies, Writing - original draft.

## Acknowledgements

The author would like to thank Xiaomeng Zhang for the coding help in finding point and cyclic attractors in this work.

## Supplementary Information

To check the accuracy of the method we tested with a slightly more connected network than the Toy model **that still allows the listing of all trajectories to each node and counting of probability distribution**. This network is Example 1 from igraph Tutorial by Chire^25^. The state transition graph is given in Figure 8. Before we can calculate the probability distribution we have to find all the trajectories to each node. We use the Python implementation from a webpage to find all paths from source to destination in a directed graph^27^. The results from our implementation are given below.

**Figure 8:**
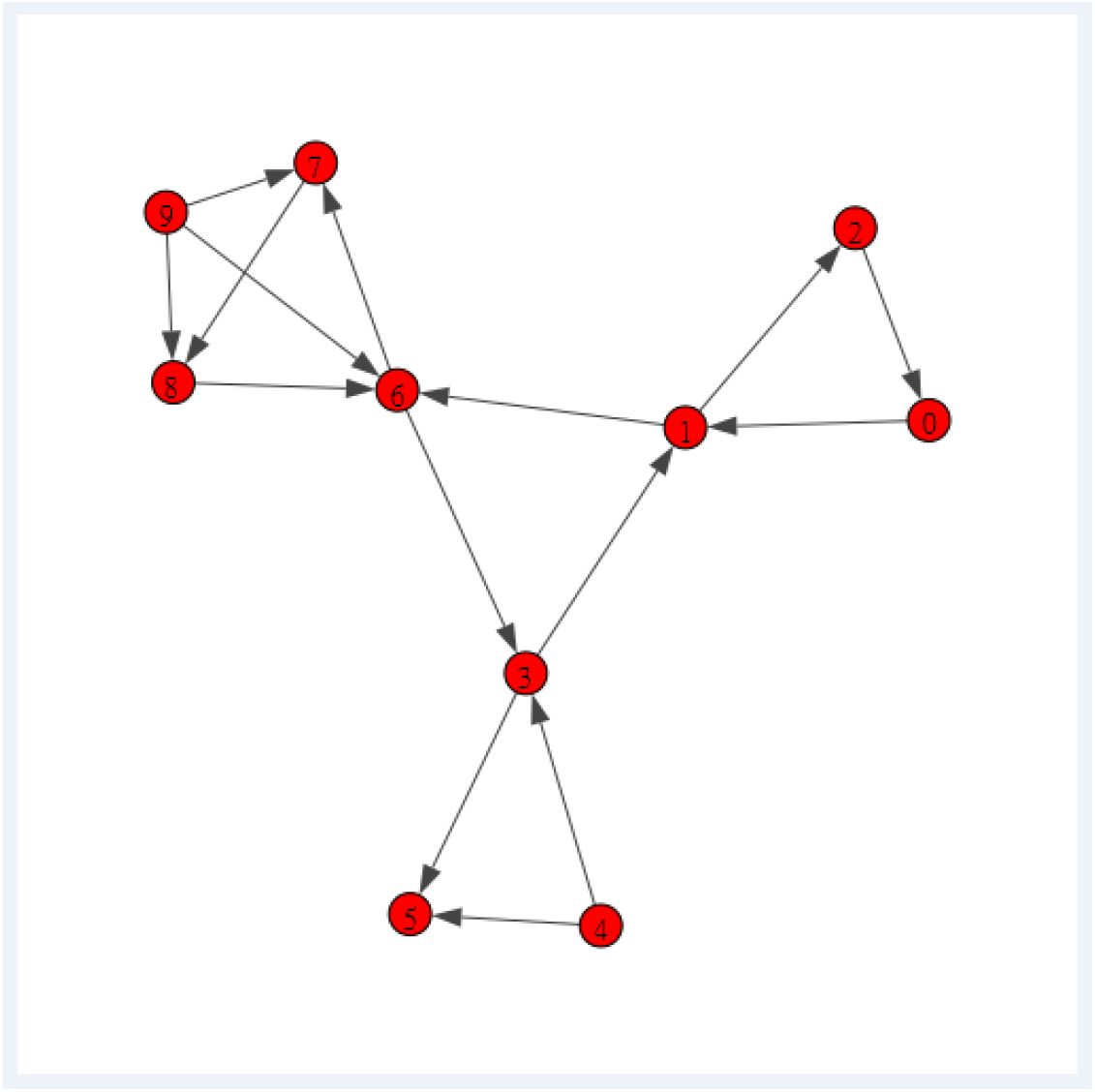
The state transition graph of a network model from Example 1 in igraph Tutorial^25^ is given above. There are 10 nodes in the network. To determine the node for cyclic attractor we need to find all cycles in a directed graph. We use the code from a webpage to find all cycles in a graph^28^. All cycles in the directed graph are: [[0, 1, 2, 0], [1, 6, 3, 1], [1, 2, 0, 1], [2, 0, 1, 2], [3, 1, 6, 3], [6, 7, 8, 6], [6, 3, 1, 6], [7, 8, 6, 7], [8, 6, 7, 8]].

The trajectories to each node are:

node [0]= [[0], [1, 2, 0], [2, 0], [3, 1, 2, 0], [4, 3, 1, 2, 0], [6, 3, 1, 2, 0], [7, 8, 6, 3, 1, 2, 0], [8, 6, 3, 1, 2, 0], [9, 6, 3, 1, 2, 0], [9, 7, 8, 6, 3, 1, 2, 0], [9, 8, 6, 3, 1, 2, 0]]

node [1]= [[0, 1], [1], [2, 0, 1], [3, 1], [4, 3, 1], [6, 3, 1], [7, 8, 6, 3, 1], [8, 6, 3, 1], [9, 6, 3, 1], [9, 7, 8, 6, 3, 1], [9, 8, 6, 3, 1]]

node [2]= [[0, 1, 2], [1, 2], [2], [3, 1, 2], [4, 3, 1, 2], [6, 3, 1, 2], [7, 8, 6, 3, 1, 2], [8, 6, 3, 1, 2], [9, 6, 3, 1, 2], [9, 7, 8, 6, 3, 1, 2], [9, 8, 6, 3, 1, 2]]

node [3]= [[0, 1, 6, 3], [1, 6, 3], [2, 0, 1, 6, 3], [3], [4, 3], [6, 3], [7, 8, 6, 3], [8, 6, 3], [9, 6, 3], [9, 7, 8, 6, 3], [9, 8, 6, 3]]

node [4]= [[4]]

node [5]= [[0, 1, 6, 3, 5], [1, 6, 3, 5], [2, 0, 1, 6, 3, 5], [3, 5], [4, 3, 5], [4, 5], [5], [6, 3, 5], [7, 8, 6, 3, 5], [8, 6, 3, 5], [9, 6, 3, 5], [9, 7, 8, 6, 3, 5], [9, 8, 6, 3, 5]]

node [6]= [[0, 1, 6], [1, 6], [2, 0, 1, 6], [3, 1, 6], [4, 3, 1, 6], [6], [7, 8, 6], [8, 6], [9, 6], [9, 7, 8, 6], [9, 8, 6]]

node [7]= [[0, 1, 6, 7], [1, 6, 7], [2, 0, 1, 6, 7], [3, 1, 6, 7], [4, 3, 1, 6, 7], [6, 7], [7], [8, 6, 7], [9, 6, 7], [9, 7], [9, 8, 6, 7]]

node [8]= [[0, 1, 6, 7, 8], [1, 6, 7, 8], [2, 0, 1, 6, 7, 8], [3, 1, 6, 7, 8], [4, 3, 1, 6, 7, 8], [6, 7, 8], [7, 8], [8], [9, 6, 7, 8], [9, 7, 8], [9, 8]]

node [9]= [[9]]

The total number of trajectories is 92. The probability distribution is given in Table 3.

**Table 3.** The probability distribution in the Chire’s igraph Tutorial Example 1 model. It is calculated based on the number of trajectory to each node listed above.

| node | no. of trajectory | probability P |
| --- | --- | --- |
| 0 | 11 | 11/92 |
| 1 | 11 | 11/92 |
| 2 | 11 | 11/92 |
| 3 | 11 | 11/92 |
| 4 | 1 | 1/92 |
| 5 | 13 | 13/92 |
| 6 | 11 | 11/92 |
| 7 | 11 | 11/92 |
| 8 | 11 | 11/92 |
| 9 | 1 | 1/92 |
| total | 92 | 1 |

The 3D landscape for the model is shown in Figure 9. The 3D landscape for Boolean network shows the attractors (1 point attractor and 7 cyclic attractors) are located at the bottom of the landscape. **The results here validated the accuracy of the implemen-tation**.

**Figure 9:**
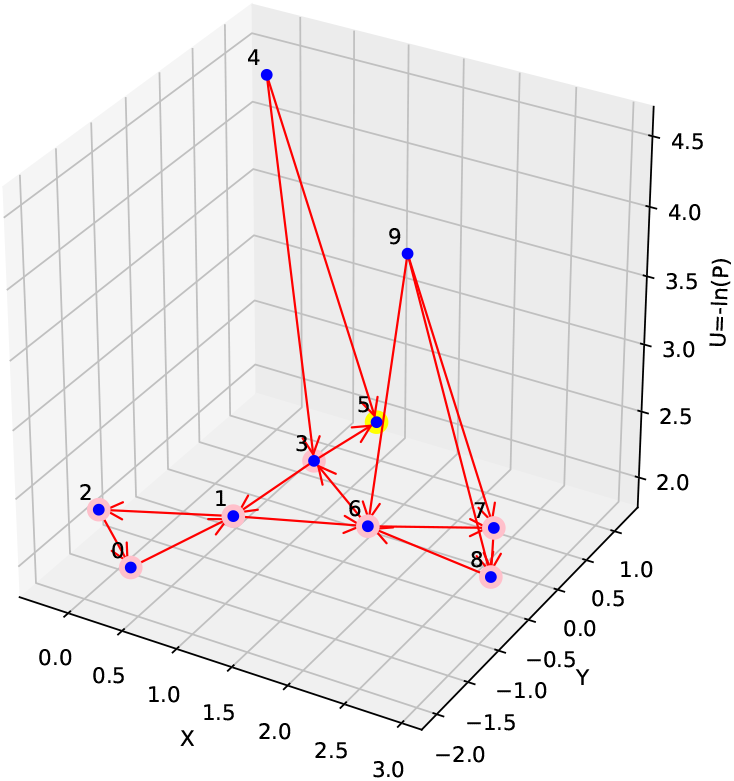
The 3D landscape of a network model from Example 1 in igraph Tutorial is given above. There is 1 point attractor (node 5 labeled yellow). Node 0, 1, 2, 6, 3, 7 and 8 are cyclic attractors (labeled pink).

